# Adaptation to developmental diet influences the response to selection on age at reproduction in the fruit fly

**DOI:** 10.1101/496836

**Authors:** Christina M. May, Joost van den Heuvel, Agnieszka Doroszuk, Katja M. Hoedjes, Thomas Flatt, Bas J. Zwaan

**Affiliations:** Laboratory of Genetics, Plant Sciences Group, Wageningen University, Wageningen, The Netherlands; Department of Ecology and Evolution, University of Lausanne, Lausanne, Switzerland; Department of Biology, University of Fribourg, Fribourg, Switzerland

## Abstract

Experimental evolution (EE) is a powerful tool for addressing how environmental factors influence life-history evolution. While in nature different selection pressures experienced across the lifespan shape life histories, EE studies typically apply selection pressures one at a time. Here we assess the consequences of adaptation to three different developmental diets in combination with classical selection for early or late reproduction in the fruit fly *Drosophila melanogaster*. We find that the response to each selection pressure is similar to that observed when they are applied independently, but the overall magnitude of the response depends on the selection regime experienced in the other life stage. For example, adaptation to increased age at reproduction increased lifespan across all diets, however, the extent of the increase was dependent on the dietary selection regime. Similarly, adaptation to a lower calorie developmental diet led to faster development and decreased adult weight, but the magnitude of the response was dependent on the age-at-reproduction selection regime. Given that multiple selection pressures are prevalent in nature, our findings suggest that trade-offs should be considered not only among traits within an organism, but also among adaptive responses to different - sometimes conflicting - selection pressures, including across life stages.

## Introduction

One of the central tenets of life-history evolution is that individuals cannot simultaneously optimize all fitness-related traits due to constraints (Roff, 1992, Roff, 2001, Stearns, 1992). These constraints can emerge because individuals have limited resources at their disposal and must make allocation decisions between competing functions (physiological constraints; Van Noordwijk & de Jong, 1986, de Jong & van Noordwijk, 1992) or because traits have a shared genetic basis (genetic constraints). Such constraints can lead to trade-offs between traits, such that an increase in one trait comes at the expense of another (Stearns, 1992).

A powerful approach for understanding how life histories and trade-offs evolve in response to specific environments is through the use of experimental evolution (EE). EE allows the experimenter to impose carefully controlled selective conditions in the laboratory and then observe evolutionary responses in real time (Kawecki et al., 2012). Two areas in which EE studies have been applied to great effect are in understanding how available nutrition influences life history evolution (Kolss et al., 2009, Kristensen et al., 2010, Leftwich et al., 2016, Chippindale et al., 1996, Bubliy & Loeschcke, 2005, Baldal et al., 2006, Zajitschek et al., 2016) and in testing the classical theories of the evolution of ageing (Luckinbill et al., 1984, Rose, 1984, Partridge & Fowler, 1992).

EE studies manipulating available nutrition have identified several correlated changes in life history traits, with the magnitude and direction of the response depending on whether the dietary manipulation is applied during development or in adulthood. Adaptation to low resource availability during development typically results in decreased adult weight (Kolss et al., 2009, Kristensen et al., 2010), faster development (Kolss et al., 2009, Leftwich et al., 2016), and lower fecundity (Kolss et al., 2009), while effects on lifespan are small or absent (Kolss et al., 2009). In contrast, adaptation to low resource availability or starvation resistance during adulthood leads to slower development, increased lipid accumulation, larger adult size, increased lifespan, and increased male fitness (Chippindale et al., 1996, Bubliy & Loeschcke, 2005, Baldal et al., 2006, Zajitschek et al., 2016, but see Hoffmann et al., 2005).

EE studies testing the classical theories of ageing have applied selection for later ages at reproduction and show that increased lifespan can reliably evolve. (Luckinbill et al., 1984, Rose, 1984, Partridge & Fowler, 1992). In most cases, decreased early or life-long fecundity is observed as a correlate of lifespan extension, suggesting a trade-off between lifespan extension and fecundity, as predicted by the disposable soma theory (Zwaan, 1999, Kirkwood & Holliday, 1979, Kirkwood & Rose, 1991).

Notably, the experiments described above each address the life history consequences of adaptation within a single life-stage and to a single selection pressure (variation in diet or selection on increased age at reproduction). However, in nature individuals will need to cope with multiple, potentially conflicting selection pressures (e.g. Lankau, 2007, Tarwater & Beissinger, 2013) experienced at different stages across the lifespan. Thus, they must balance the relative costs and benefits of adaptation and resource allocation made at one life stage with those at other stages (reviewed in Schluter *et al*., 1991). Indeed, EE studies applying more than one selection pressure within a single life stage reveal that the responses to multiple selection pressures tend to be interdependent (Davidowitz et al., 2016, Bochdanovits & Jong, 2003), yet also - despite constraining correlations among traits - there is potential for independent evolutionary change (Beldade & Brakefield, 2002, Frankino et al., 2005). To date, however, there has been little emphasis on how multiple selection pressures influence life histories as a whole. In particular, to the best of our knowledge, no study to date has combined two selection regimes experienced at different stages across an organism’s lifespan.

Here we combine variation in available nutrition during development with classical selection for early or late reproduction during adulthood in a single fully-factorial EE design, using the fruit fly, *Drosophila melanogaster* (Fig. 1a). Empirical work suggests that the two selection regimes might exert opposing selection pressures, which will have to be integrated into the life history. For example, adaptation to a poor quality diet generally selects for faster development coupled with smaller adult size and decreased fecundity (Bochdanovits & Jong, 2003, Kolss et al., 2009), whereas longer lifespan (the typical response to selection on increased age at reproduction) is generally correlated with longer developmental time and larger size (Lints, 1978, Economos, 1980, Promislow, 1993, Khazaeli et al., 2005, but see Zwaan et al., 1991).

**Figure 1.**
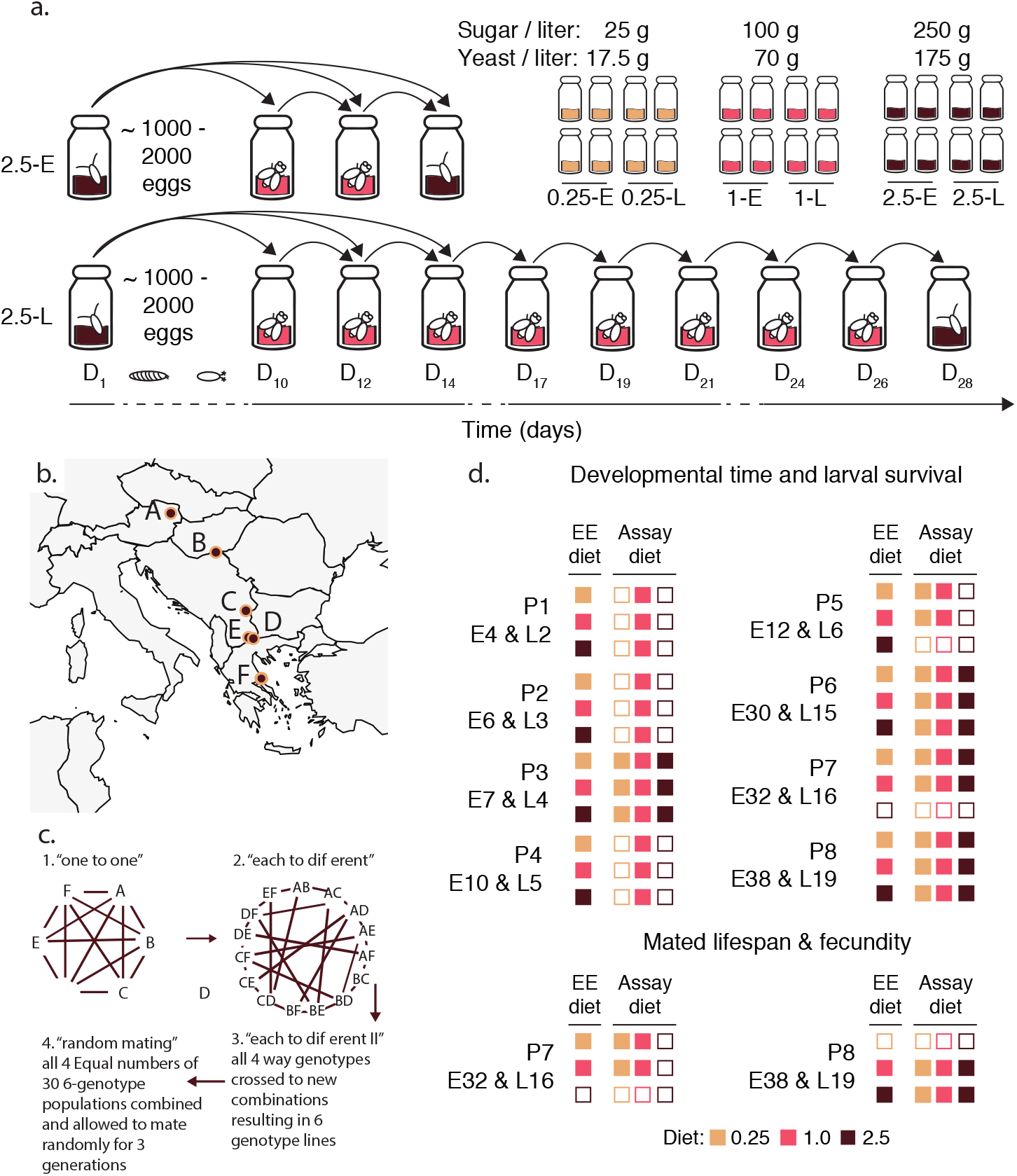
Experimental overview. (a) Experimental evolution design. Four replicate populations were established per combination of larval diet and age at reproduction (4 replicate lines * 3 larval diets * 2 ages at reproduction = 24 lines in total). The main panel traces a generation of EE for a single 2.5SY-E (top) and 2.5SY-L (bottom) line. (b) Collection sites across Europe of the six populations that contributed to the “S” starting population. (c) A brief description of the multi-generation crossing scheme used to cross the six populations in (b) to generate the mixed “S” population used for experimental evolution. (d) Overview of traits assayed in each phenotyping session (8 in total, labeled P1 through P8). Inclusion of lines and assay diets in a phenotyping is diagrammed using filled (included) vs. unfilled boxes (not included).. Briefly, the first column of boxes indicates the experimental evolution lines included, while the second, third and fourth columns indicate the assay diets on which they were assessed (key in inset box). In all cases, both the early (E) and late (L) reproducing lines were included. Thus, for example, in P4 (Generation 10 & 5 of E and L lines respectively) the 0.25-E, 0.25-L, 1-E, 1-L, 2.5-E, and 2.5-L lines were all included (first column all filled), but only assayed under the 1.0 assay diet. It is noteworthy that there is a relatively large generation gap between sessions P1 through P5 and P6 through P8.

Our experimental design allows us to address several fundamental problems, including the question of whether adaptation to environmental variation in each stage occurs independently. For example, will a lower calorie developmental diet constrain the ability to extend lifespan in response to selection on age at reproduction, or will lifespan extension be achieved at the expense of other traits? To address this issue we assess the evolutionary responses of several life-history traits. These include larval survival, developmental time, and adult weight, all of which have previously been found to evolve in response to larval acquisition (e.g. Bochdanovits & Jong, 2003, Kolss et al., 2009), as well as adult lifespan and fecundity, the two traits that commonly trade off in response to selection on age at reproduction (Luckinbill et al., 1984, Rose, 1984). Furthermore, we assay traits over multiple generations and in multiple larval dietary environments to gain insight into the temporal dynamics of evolution, and the evolution of phenotypic plasticity.

## Materials and Methods

### Design of the experimental evolution experiment

We combined three levels of larval diet (0.25, 1.0 and 2.5) with two ages at reproduction (Early and Late) in a fully factorial design (Fig.1a, inset). The larval diets differed only in sugar and yeast content with the 0.25 and 2.5 diets containing 25% and 250% as much sugar and yeast as the 1.0 diet, respectively (Table S1). These diets were chosen to fall within the range typically applied in studies of diet and life history in *D. melanogaster* (Zajitschek et al., 2016, Magwere et al., 2004, Lee et al., 2008). The early (E) and late (L) reproducing populations had generation times of 14 and 28 days respectively, thus adults laid eggs for the subsequent generation roughly two to four days post-eclosion in the Early (E) lines, and 16 to 18 days post-eclosion in the Late (L) lines (Fig. 1a). For each combination of larval diet and age at reproduction, we established four independent replicate lines (3 larval diets x 2 ages at reproduction x 4 replicate lines = 24 lines total; Fig. 1a, inset). All lines were maintained on the 1.0 diet as adults both in the course of evolution and in all experiments. We refer to the EE lines by their larval diet (0.25, 1.0, 2.5), their age at reproduction (E, L), or the combination of the two (0.25-E, 0.25-L, 1-E, 1-L, 2.5-E and 2.5-L) throughout. Since the diet and age-at-reproduction conditions of the 1-E lines mimic those of our standard laboratory maintenance regime, their responses can be considered representative of the baseline response both in terms of plasticity and the evolutionary response across generations. Lines were maintained throughout under standard laboratory conditions (25°C, 65% relative air humidity, 12 h: 12 h light : dark cycle).

### Generating the starting population and initiating experimental evolution

To ensure ample standing genetic variation the EE populations were derived from six populations of flies collected along a latitudinal gradient across Europe (Fig. 1b). These populations were maintained in the laboratory for 40 generations to allow for laboratory adaptation and then combined into a single panmictic, genetically diverse baseline population, the starting (“S”) population, using a multi-generation crossing scheme (Fig. 1c; see May et al., 2015 for full details of the crossing scheme). This scheme was employed to minimize linkage disequilibrium and to ensure equal contributions of the component populations to the final “S” population. After crossing, the “S” population was maintained under standard laboratory conditions for a further 10 generations at a population size of ~ 4000 individuals.

To initiate EE, eggs were collected from the “S” population into large glass bottles (500 mL volume) filled with 65 mL of the respective larval diets. Two bottles of ~ 1000-2000 eggs were collected per replicate line and allowed to develop to adulthood. For each larval diet four lines were randomly assigned to the early (E) and late (L) reproduction regimes. Ten days after egg laying (Monday) we transferred all newly eclosed adults into fresh bottles containing the 1.0 diet. Populations were then transferred to fresh bottles of 1.0 medium every Monday, Wednesday, and Friday until their respective ages at reproduction. Since larval diet affected developmental time, not all adults from all lines had emerged by day 10, so on days 12 and 14 any additional late-eclosing adults from the developmental bottles were added to the adult population bottles to mitigate truncation selection on developmental time (Fig. 1a). Very few flies eclosed after day 12.

The day before populations reached their respective ages at reproduction, 1/16 of a teaspoon of dry yeast (Fermipan Red Instant dry bakers yeast) was added to each bottle to stimulate females to lay eggs. The following day females were transferred to fresh bottles containing their evolutionary larval diets and allowed to lay eggs. A test-tube cap containing dry yeast mixed with water was suspended in the bottle and removed when egg laying was complete so as not to modify yeast levels in the developmental diet. To control egg densities, bottles were visually inspected, and adult flies were removed from bottles when egg density was between ~ 1000-2000 eggs, typically over a period of two to four hours. Every generation, both replicate bottles within a line were mixed. Overall, population size was ~ 2000 to 4000 adult flies per replicate line over the course of EE.

### Assessing changes in life history traits over the course of evolution

We measured four key life-history traits: egg-to-adult development time, mated female fecundity, mated lifespan and adult wet weight. We assayed these traits across eight independent phenotyping sessions, ranging from the beginning of EE up to generations 38 and 19 for E and L lines, respectively. Figure 1d provides an overview of each phenotyping session (P1-P8), including the elapsed generations of EE, the lines included, the traits measured, and the larval conditions under which flies were raised (i.e., assay environment). We deliberately chose larval diets that had negligible effects on larval survival to avoid population bottlenecks and strong viability selection. Larval survival ranged from 80-95% across evolutionary larval diets and assay conditions in all but one phenotyping session (Supplementary Figure 1) and did not show any systematic variation across selection regimes (Table 2 and Supplementary Figure 1), suggesting that larval survival was not under selection.

Whenever possible we measured the responses to selection in all lines and used all three larval assay diets. However, the scale of our design imposed some logistical constraints. In some phenotyping sessions, we monitored the progress of adaptation on the 1.0 larval assay diet only, while in others we raised larvae on all three diets. In all cases, we first allowed lines to develop for one generation on the 1.0 diet to avoid potential maternal effects. Larvae developed at a density of 70 eggs per vial, with 6 mL of food per vial. For each line, eggs were collected from petridishes and randomized across assay diets.

We assessed development time and survival from egg to adult in all eight phenotyping sessions (Fig. 1d; n=5 vials per combination of line and assay diet). We scored developmental time until no new flies emerged over a period of 48 hours and then summed across the resulting adults to obtain a measure of egg-to-adult survival (proportion viability). While using vials allowed easier standardization of egg densities and more accurate counting of eclosing adults, development took ~ 24 hours longer in vials than in the EE population bottles.

Mated lifespan and fecundity were assessed on the evolutionary larval diet and on the 1.0 diet The size of this experiment necessitated two assays (Fig. 1d): in the first round, we tested all lines adapted to 0.25 or 1.0 larval food on these two diets (P7), and in the second round we tested all lines adapted to 1.0 and 2.5 larval diet across all three larval diets (P8). The 1.0 lines served as a reference to facilitate comparisons between the 0.25 and 2.5 lines and to monitor consistency of responses across both assays. For mated lifespan, we housed flies at a density of three males and three females per vial (n=10 vials per combination of line and larval diet). Flies were transferred to fresh vials and survival was scored every Monday, Wednesday, and Friday.

Mated fecundity was measured over three time spans: Early (days 2-4 of adulthood), Late (days 18-21) and Post-selection (days 25-28) with the Early and Late time points overlapping the ages at reproduction of the E and L lines. In the first assay (P7), we maintained a single male-female pair per vial (n=15 vials per line and larval food combination), while in the second assay (P8) we maintained two males and two females per vial (n=10 vials per line and larval food combination). Eggs were allowed to develop to adulthood and emerging adults were counted to score fecundity. Sperm depletion was prevented by replacing dead males with new males from the same experimental conditions.

Wet weight of adult males and females raised on the 1.0 assay diet was obtained in generations 144 and 73 of the E and L lines respectively. All 24 EE populations were reared in small bottles (200 mL) with 25 mL 1.0 food at a density of 600-800 eggs per bottle (Clancy & Kennington, 2001). After eclosion, males and females were housed together until weighing (i.e., they were mated). Weight was measured at two time points chosen to mimic the conditions of the EE procedure: 14 days after egg laying (~4-5 days after emergence) and 28 days after egg laying (~18-19 days after emergence). The weight of the flies was measured on an ultramicro balance (Sartorius Cubis Ultramicro Balance MSE) using batches consisting of two flies each (n=10). Prior to the assay, all populations were first reared for two generations on 1.0 medium. This assay was performed later than the other life-history assays, however, the results were consistent with an interim measurement made on a subset of the lines at ER generation 100 and LR generation 50 (data not shown).

### Statistical analysis

All statistical analyses were performed in R version 3.2.0 (R Development Core Team, 2005). We fitted a separate model for each trait within each phenotyping session. In each model we included evolutionary dietary regime, age at reproduction, assay diet, sex (where applicable), and their interactions as explanatory variables. We used mixed-effects models to accommodate the random effect of replicate line nested within a selection regime. Both developmental time and mated longevity were analyzed using mixed-effects Cox regression (proportional hazards) models (coxme package; Therneau, 2015), while larval survival and fecundity were analyzed with generalized linear mixed models (GLMM) with binomial and Poisson error distributions, respectively (lme4 package; Bates et al., 2015). Weight was analyzed using a linear mixed-effects model with a normal distribution. In the statistical tables (see below), we report the *χ*^2^values of the effect of each factor in the full model as obtained by Analysis of Deviance (car package; Fox & Weisberg, 2010). We performed further model simplification by sequentially dropping non-significant terms from the model and using a *χ*^2^ test to compare models. To control for multiple comparisons we applied the sequential Holm-Bonferroni correction method to each fitted model (Holm, 1979) and the Tukey’s range test for all post-hoc comparisons among means (Tukey, 1949).

For each trait assessed across multiple assay diets we fitted an additional model for the 1-E lines alone (unselected control lines) to determine the baseline plastic response. An inconsistent response of the 1-E lines across generations might indicate that the trait in question is sensitive to slight differences in developmental conditions; in this case, differences among lines might be highly dependent on variations in assay conditions and may thus not reflect robust evolutionary responses (cf. Ackermann et al. 2001).

## Results

### Developmental time depends on selection regime by assay diet interactions

Over the course of evolution, assay diet was the most important factor influencing developmental time (Table 1; Fig. 2). The 0.25, and to a lesser extent, 2.5 assay diet consistently increased developmental time relative to the 1.0 diet across all EE lines and phenotyping sessions (Fig. 2 p<0.00001 in all but one contrast). However, it is noteworthy that the mean duration of development on each of the three assay diets fluctuated greatly across phenotyping sessions (Fig. 2). Such variation is not uncommon in repeated measures of developmental time (e.g. Zwaan et al., 1995). To account for this, we plotted both the absolute values of developmental time per assay diet across phenotyping assays (Fig. 3 a-c) and relative to the mean of the 1-E lines (Fig. 3d-f).

**Table 1:** Summary of GLMMs (Chi-square values) for the effect of assay diet (A), evolutionary dietary regime (D) and evolutionary age at reproduction (R) on larval survival and developmental time across phenotyping sessions. Significance is indicated by: * =P<0.05, **=P<0.01, ***=P<0.001.

| Phenotyping | EE Diet (D) | EE Repro (R) | Assay diet (A) | D*R | D*A | R*A | D*R*A |
| --- | --- | --- | --- | --- | --- | --- | --- |
| **Larval survival** |  |  |  |  |  |  |  |
| P1 | 0.26 | 0.05 | --- | 0.24 | --- | --- | --- |
| P2 | 0.30 | 0.85 | --- | 0.37 | --- | --- | --- |
| P3 | 0.71 | 0.02 | 6.6 | 0.57 | 1.50 | 0.00 | 0.66 |
| P4 | 0.35 | 0.01 | 1.24 | --- | --- | --- | --- |
| P5 | 13.06** | 3.94 | 0.17 | 0.32 | 25.24*** | 0.46 | 0.85 |
| P6 | 5.23 | 2.49 | 42.69*** | 2.73 | 5.99 | 38.86*** | 12.91 |
| P7 | 0.00 | 0.00 | 3.23 | 0.12 | 3.02 | 0.74 | 0.51 |
| P8 | 1.45 | 0.04 | 8.07 | 0.62 | 2.68 | 0.43 | 3.66 |
| **Developmental time** |  |  |  |  |  |  |  |
| P1 | 2.07 | 27.58*** | --- | 0.96 | --- | --- | --- |
| P2 | 4.14 | 0.63 | --- | 0.68 | --- | --- | --- |
| P3 | 0.46 | 15.74*** | 20617.4*** | 0.34 | 140.61*** | 64.27*** | 72.31*** |
| P4 | 3.86 | 2 | ---- | 0.97 | ---- | ---- | ---- |
| P5 | 0.18 | 4.30 | 4830.06*** | 2.44 | 25.13*** | 244.81*** | 41.63*** |
| P6 | 8.06 | 2.02 | 12746.57*** | 11.57**<br>1.06 | 76.11*** | 311.47*** | 48.51*** |
| P7 | 6.90 | 0.18 | 8506.83***<br>23285*** |  | 22.29*** | 34.18*** | 78.39*** |
| P8 | 7.64 | 13.36*** |  | 1.21 | 181.40*** | 758.70*** | 46.53*** |

**Figure 2.**
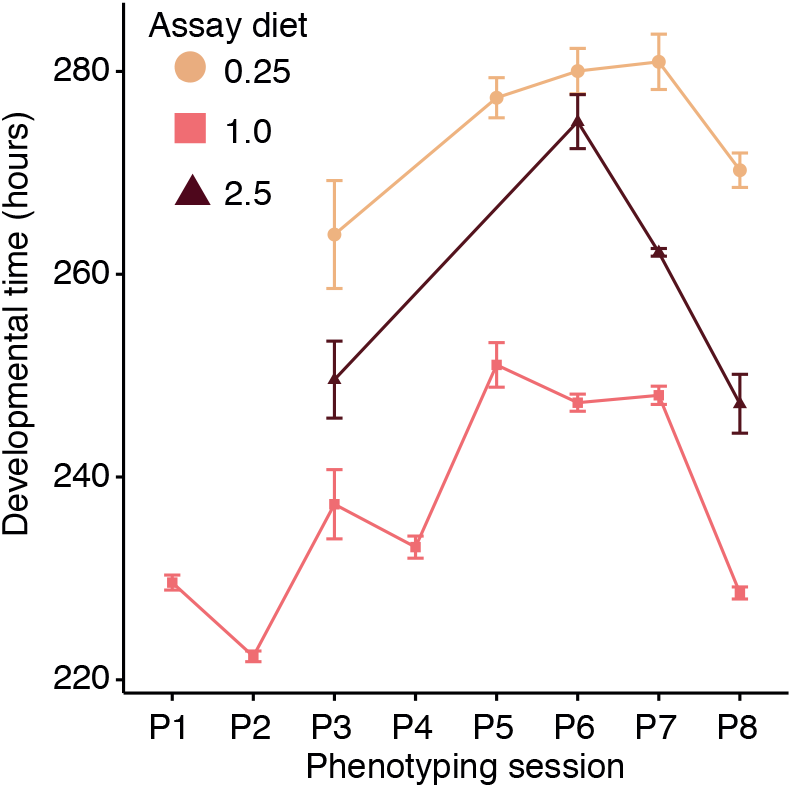
Developmental time from egg to adult (y-axis) across phenotyping sessions (x-axis) by assay diet (0.25: beige square; 1.0 : pink circle; 2.5: purple triangle). Not all phenotyping sessions included all three assay diets. Each point represents taking the mean of the average developmental time for each of the lines included in the assay. For example, if all 24 lines were included in the assay the mean developmental time was calculated per line and then the mean and standard error of these 24 values was calculated.

**Figure 3.**
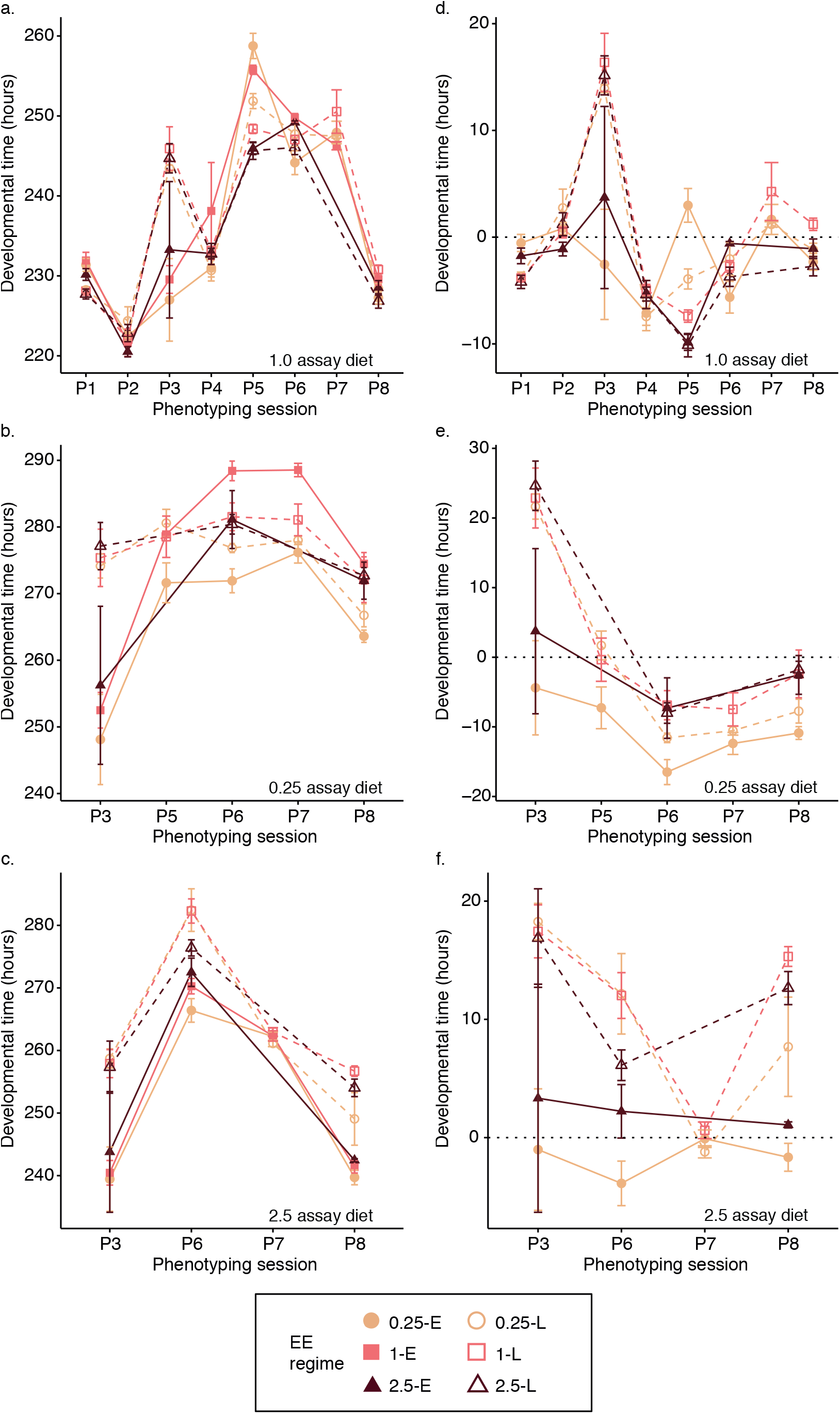
Developmental time from egg to adult (y-axis) across phenotyping sessions (x-axis) for the 1.0 assay diet (a,d), the 0.25 assay diet (b,e) and the 2.5 assay diet (c,f). (a-c) represent the observed developmental times while (d-f) are the developmental times relative to the mean of the 1-E lines. Each point represents taking the mean and standard error of the average developmental time for each of the four lines.

In the early generations of EE (P1-P5) no consistent changes in developmental time were observed (Fig. 3). However, from P6 (E and L generations 30 and 15, respectively) onwards a consistent three-way interaction emerged between evolutionary larval diet, age at reproduction, and assay diet for the 0.25-E and 0.25-L lines (Table 1; Fig. 3). Both sets of lines evolved substantially more rapid development on the 0.25 assay diet as compared to the 1-E lines (Fig. 3b,d). For the 0.25-E lines this effect was already present in P5 (P5 through to P8; all p-values <0.001), while for the 0.25-L lines it became apparent from P6 onwards (P6 through to P8; all p-values <0.001; Fig. 3b,e), although there was a trend for the magnitude of the effect to be smaller for the 0.25-L than 0.25-E lines (P5: p<0.0001; P6: p=<0.01; P7: p=0.33; P8: p=0.26). By contrast, on the 1.0 and 2.5 assay diets the responses of the 0.25-E and 0.25-L lines were not consistent across phenotypings (Fig. 3a,c,d,f). Relative to the strength and consistency of the response of the 0.25-E and 0.25-L lines, the 1-L and 2.5-E and 2.5-L lines did not show substantial or consistent changes in developmental time, suggesting that these regimes did not impose strong selection on the length of development.

### Selection on age at reproduction increases lifespan across dietary selection regimes

We found that selection for increased age at reproduction increased lifespan in all lines and across all assay diets (Fig. 4; all p-values <0.03). However, the magnitude of the effect was dependent on sex and the evolutionary dietary regime for 0.25 lines and evolutionary dietary regime and assay diet for 2.5 lines, suggesting that adaptation to different levels of laerval acquisition can modify the response to selection on lifespan (Table 3).

**Figure 4.**
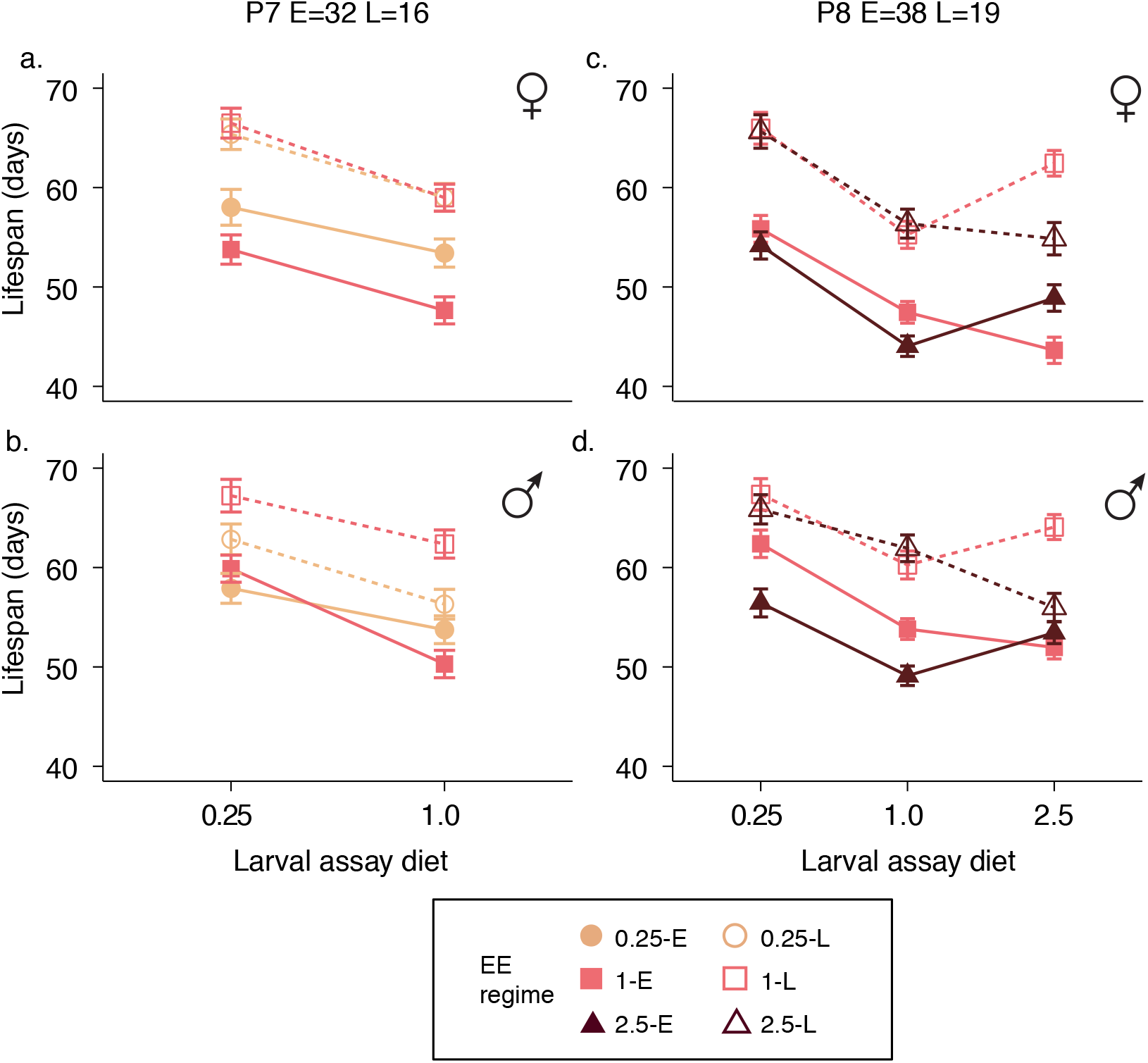
Lifespan (y-axis) across assay diets (x-axis) and phenotyping sessions P7 (a, b) and P8 (c, d) for females (a, c) and males (b, d). Lifespan is expressed as days from adult eclosion. All error bars are standard errors of the mean across replicate lines.

For the 0.25-E and 0.25-L lines males and females had inverse responses to selection relative to the 1-E and 1-L lines (Table 3; Fig. 4a,b). In females, the lifespan of the 0.25-L lines was indistinguishable from that of 1-L lines (p=0.94), but 0.25-E lines had greater lifespans relative to the 1-E lines (p<0.0001) (Fig. 4a). In males, the exact inverse response was observed: while the 0.25-E and 1-E lines had similar lifespans (p=0.96), the lifespan extension of 0.25-L lines was less than that of the 1-L lines (p=0.01; Fig. 4b). These effects were consistent across both the 0.25 and 1.0 assay diets.

For the 2.5-E and 2.5-L lines, lifespan evolved in a similar manner in both sexes, but was dependent on assay diet (Table 3 and Figure 4c,d). Under 0.25 and 1.0 assay conditions, the lifespan of the 2.5L lines did not differ from the 1-L lines (add p-value for both sexes), however, under 2.5 assay conditions 2.5-L flies evolved significantly shorter lifespans than 1-L lines in males (p=0.002, Fig. 4d) and nearly significantly shorter lifespans in females (p=0.08, Fig. 4c). The 2.5E lines showed an inverse pattern: lifespan on the 0.25 and 1.0 assay diets was generally higher for 1-E lines than for 2.5-E lines, whereas males and females of the 2.5-E lines outlived 1-E flies on the 2.5 assay diet (Fig. 4c,d; Males: on 0.25 and 1.0 diet 1-E>2.5-E, p=0.003 and 0.004, respectively; under 2.5 assay conditions 1-E=2.5-E, p=0.66. Females: on 0.25 assay diet 1-E=2.5-E, p=0.42; on 1.0 assay diet 1-E>2.5-E lines, p=0.02; and under 2.5 assay diet 1-E<2.5-E lines, p=0.01).

### Fecundity is highly variable across phenotyping sessions

Because it was not possible to measure lifespan and fecundity for all lines at the same time, we used the 1-E and 1-L lines as a standard across the two replicate phenotyping sessions (see Materials and Methods). For mated fecundity, the plastic response of the 1-E lines to assay diet differed between the two phenotyping sessions (Table 2, Fig. 5). In the first phenotyping (P7) 1-E flies raised on the 0.25 assay diet had lower fecundity than those raised on the 1.0 assay diet at all three ages (Fig. 5a-c; all p-values<0.001). In the second assay (P8), the same effect was observed at early and post-selection ages (all p-values<0.001), but reversed at the late reproduction time point (p<0.001; Fig 5d, f). Furthermore, the difference between the 1-E and 1-L lines on the 1.0 assay diet was also inconsistent between assays P7 and P8 (Fig. 6). In P7 the E lines reproduced more than the L lines at the “Mid” time point and less at the “Late” time point, while in P8 the opposite pattern was observed (Fig.6, both p-values <0.003). Thus, while the GLMM’s indicated that fecundity at all ages was affected by interactions between diet regime, age at reproduction regime, and assay conditions (Table 4), the lack of consistency of the 1-E and 1-L lines hampers the interpretation of the evolutionary significance of these effects.

**Table 2:** Summary of GLMMs (Chi-square statistics, degrees of freedom, and their significance) for the effect of assay diet on larval survival, developmental time, lifespan, and fecundity on 1-E lines across phenotyping sessions. Where there was a significant effect of assay diet (i.e. plasticity for the response to assay diet) we report the outcomes of pairwise post-hoc comparisons between assay diets (p-values). Where several models were fit per trait we indicate the subset analyzed (Subset).

| Phenotyping | Generation | Subset | Effect Assay diet |  |  | Post-hoc contrasts |  |  |
| --- | --- | --- | --- | --- | --- | --- | --- | --- |
|  |  |  | Chi square | df | p-value | P:C | P:R | R:C |
| Larval survival |  |  |  |  |  |  |  |  |
| P3 | 7 | --- | 1.59 | 2 | 0.45 | --- | --- | --- |
| P5 | 12 | --- | 6.18 | 1 | 0.01 | 0.01 | --- | --- |
| P6 | 30 | --- | 0.42 | 2 | 0.81 | --- | --- | --- |
| P7 | 32 | --- | 0.44 | 2 | 0.80 | --- | --- | --- |
| P8 | 38 | --- | 5.46 | 2 | 0.07 | --- | --- | --- |
| Developmental time |  |  |  |  |  |  |  |  |
| P3 | 7 | --- | 1878.70 | 2 | <0.0001 | <0.0001 | <0.0001 | <0.0001 |
| P5 | 12 | --- | 1090.70 | 1 | <0.0001 | <0.0001 | --- | --- |
| P6 | 30 | --- | 2648.30 | 2 | <0.0001 | <0.0001 | <0.0001 | <0.0001 |
| P7 | 32 | --- | 2303.30 | 2 | <0.0001 | <0.0001 | <0.0001 | <0.0001 |
| P8 | 38 | --- | 4212.50 | 2 | <0.0001 | <0.0001 | <0.0001 | <0.0001 |
| Fecundity |  |  |  |  |  |  |  |  |
| P7 | 32 | Early | 12.904 | 1 | <0.0001 | 0.0001 | --- | --- |
|  |  | Mid | 570.12 | 1 | <0.0001 | <0.0001 | --- | --- |
|  |  | Late | 392.35 | 1 | <0.0001 | <0.0001 | --- | --- |
| P8 | 38 | Early | 251.46 | 2 | <0.0001 | <0.0001 | <0.0001 | 0.79 |
|  |  | Mid | 225.24 | 2 | <0.0001 | <0.0001 | <0.0001 | <0.0001 |
|  |  | Late | 65.824 | 2 | <0.0001 | <0.0001 | 0.42 | <0.0001 |
| Lifespan |  |  |  |  |  |  |  |  |
| P7 | 32 | F | 16.015 | 1 | <0.0001 | <0.0001 | --- | --- |
|  |  | M | 32.831 | 1 | <0.0001 | <0.0001 | --- | --- |
| P8 | 38 | F | 55.467 | 2 | <0.0001 | <0.0001 | <0.0001 | 0.23 |
|  |  | M | 46.134 | 2 | <0.0001 | <0.0001 | <0.0001 | 0.45 |

**Figure 5.**
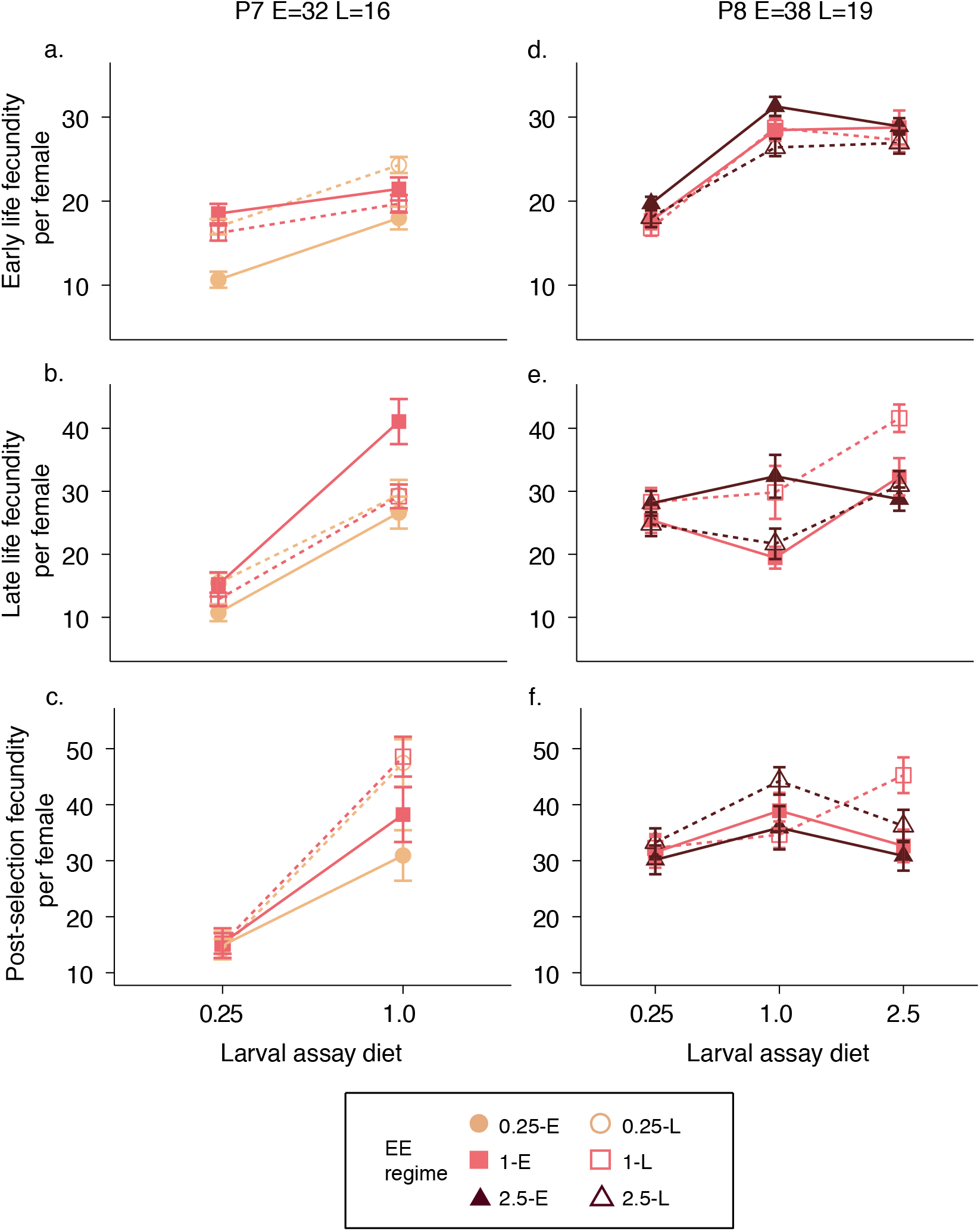
Reaction norms of realized early (a,d), late (b,e) and post-selection (c,f) female fecundity (y-axis) across assay diets (x-axis) and phenotyping sessions P7 (a:c) and P8 (d:f). All error bars are standard errors of the mean across replicate lines.

**Figure 6.**
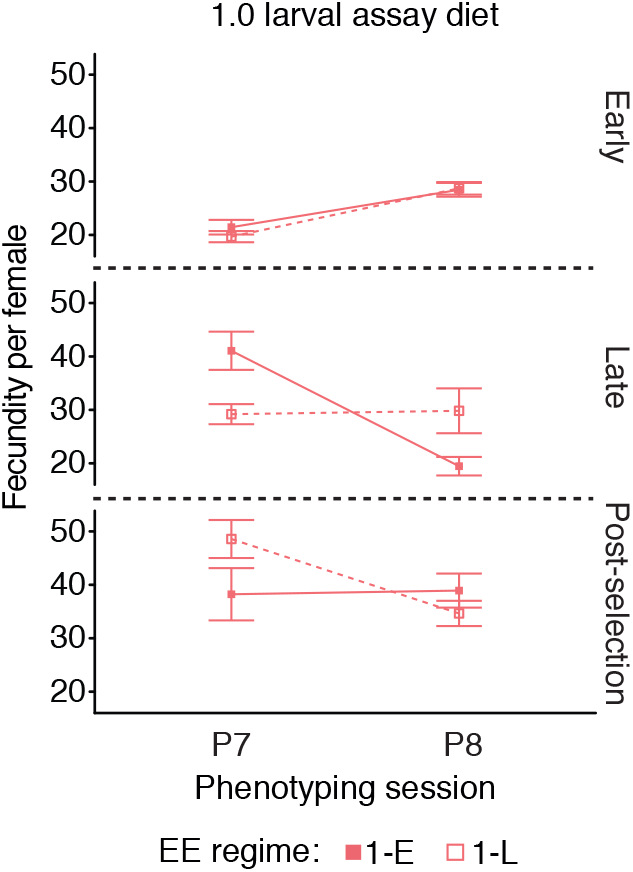
Inconsistencies in fecundity of 1.0 lines across phenotyping sessions. All error bars are standard errors of the mean across replicate lines.

### Adult weight

Adult weight evolved in response to the selection regimes in a sex- and age-dependent manner (Fig. 7a-d). The largest effects of the EE regimes occurred in young flies (4-5-days post-eclosion): in both sexes adaptation to later ages-at-reproduction led to larger adult size (Females: F_1,24_=8.4, *p* =0.01; Fig. 7a; Males: F_1,24_=43.7, *p*=<0.0001; Fig.7b), and adaptation to the 0.25 larval diet decreased adult weight relative to 1.0 and 2.5 adapted lines (Females:F_2,24_=10.9, *p* =0.001; Fig. 7a; Males: F_2,24_=29.6, *p*=<0.0001; Fig.7b; all pair-wise p-values <0.01). In females, there was a marginal interaction between age at reproduction and evolutionary dietary regime (F_2,24_=2.7, *p* =0.09, Fig. 7a). While the 0.25-L and 1-L lines both evolved increased weights relative to the 0.25-E and 1-E lines, the weight of the 2.5-E and 2.5-L lines did not differ (2.5-L = 2.5-E, p=1.0). At 18-19 days post-eclosion the effects of evolutionary regime became much smaller and differed between the sexes. In males, the effect of EE regime was largely absent, except in 0.25-E lines, which continued to weigh less than all other lines (all pairwise p-values <0.003; Fig. 7d), while in females, only evolutionary dietary regime remained significant (F_2,24_=9.1, p=0.001, Fig. 7c), with weight increasing with increasing evolutionary larval diet (all pairwise p-values <0.05). We also found large effects that were independent of the evolutionary regimes: males weighed less than females (Sex: F_1,936_=11644, *p*=<0.0001) and, while females gained weight with age, males tended to lose or maintain the same weight (Sex x Age: F_1,936_=314.3, *p*=<0.0001; Fig.7).

**Figure 7.**
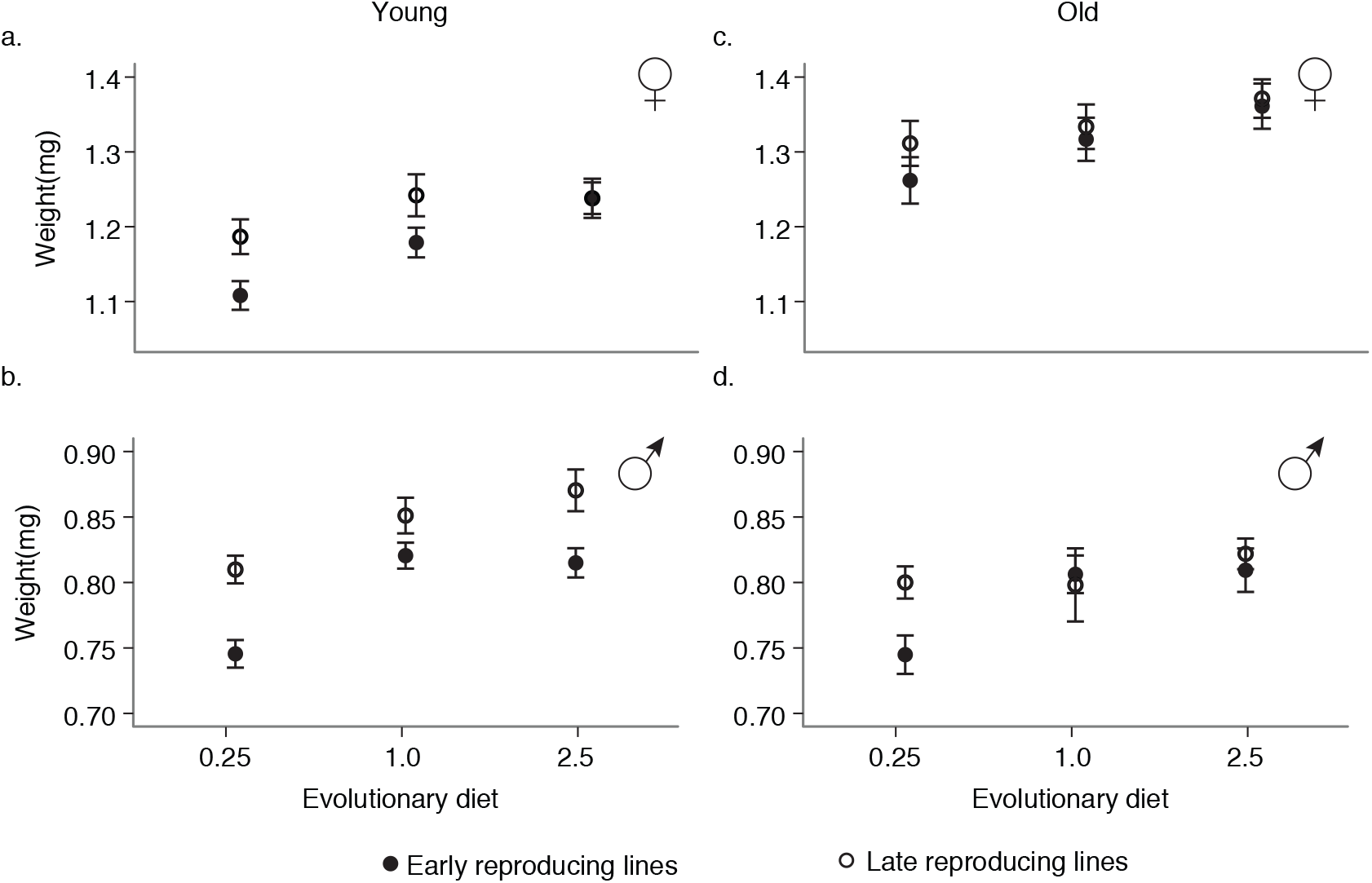
Body weight (mg) of female (a,c) and male (b,d) flies raised on the 1.0 assay diet at young (~ 4-5 days old) and old (~18-19 days old) ages. All error bars are standard errors of the mean across replicate lines.

**Figure 8.**
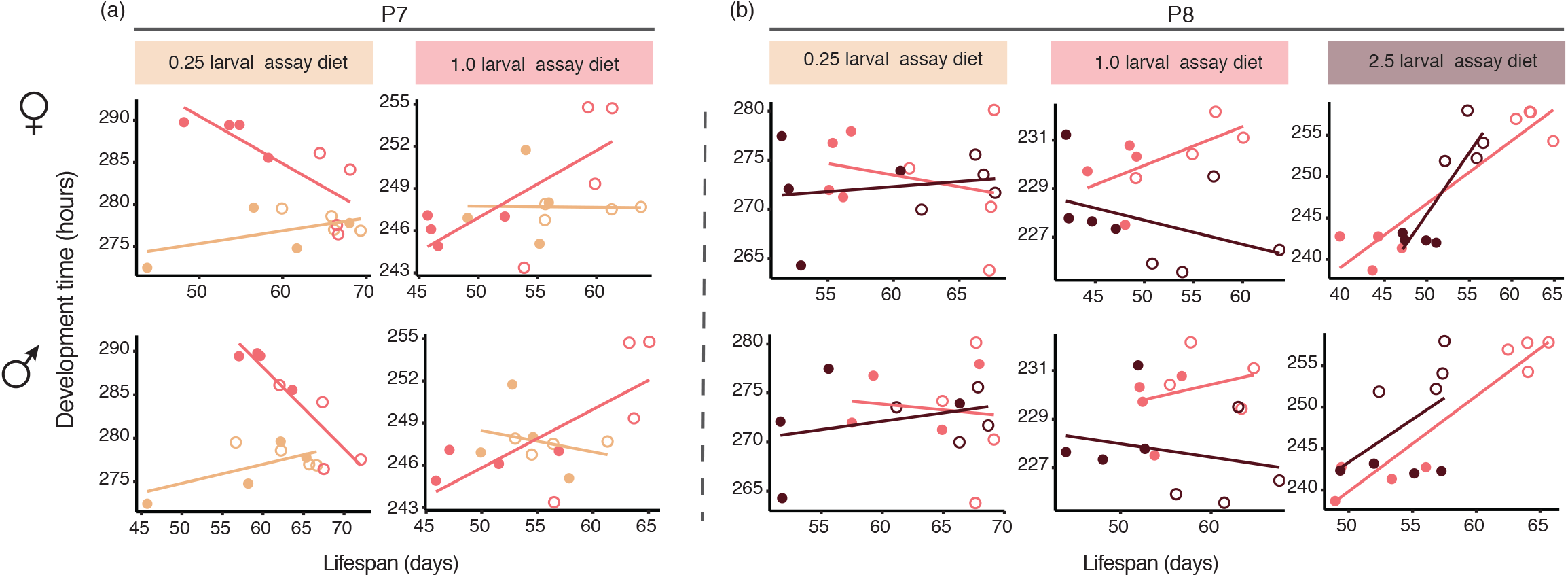

## Discussion

How different selection pressures interact to affect life-history adaptation is an unresolved question. By utilizing the combined strength of extensive replication, multiple assay environments, and assessment of evolution across multiple generations we were able to discriminate between transient and consistent effects of adaptation to larval diet and age at reproduction. We discuss our main findings in the light of theoretical predictions and previous work.

### Adaptive responses reflect the influence of both selection regimes

For both selection regimes we observed similar changes in life history traits to those observed in previous, univariate studies. That is, adaptation to increased age at reproduction increased lifespan across all evolutionary dietary regimes and in both sexes (Luckinbill et al., 1984, Rose, 1984, Partridge & Fowler, 1992), while selection on the 0.25 larval diet resulted in faster development (Fig. 3b,e), decreased adult weight (Fig. 7), and potentially lower fecundity (Fig.5a-c), again, in keeping with previous univariate selection experiments (Kristensen et al., 2010, Kolss et al., 2009). However, in both cases, we found that the addition of a second regime modified the magnitude of the responses. Thus the extent of the increase in lifespan imposed by selection for later age at reproduction was dependent on dietary regime (Fig. 4) and conversely, the changes in weight, length of development and potentially fecundity seen in the 0.25-E lines were modified by adding selection for late reproduction. Thus, while the two regimes continue to select for similar adaptive responses, the overall magnitude of the response depends on the interplay with the selection pressure experienced in the other life stage.

### Fecundity: significant but inconsistent responses

Previous EE designs selecting on later age at reproduction also found inconsistent responses of fecundity across generations (Leroi et al., 1994a), or marked sensitivity to environmental variation (Leroi et al., 1994b). However, we observed strongly significant effects of both age at reproduction and evolutionary dietary regime in both phenotyping sessions (Table 4). For example, 0.25-E lines appeared to have decreased fecundity relative to 0.25-L, 1-E and 1-L lines at all ages (Fig. 5a-c), a response that is consistent with their lower body weight and faster development (Fig. 3b,e and Fig. 7). Given the large replication of our design (i.e., independent replicate populations per EE treatment) it is plausible that these responses represent adaptive responses to poor nutrition.

**Table 3:**
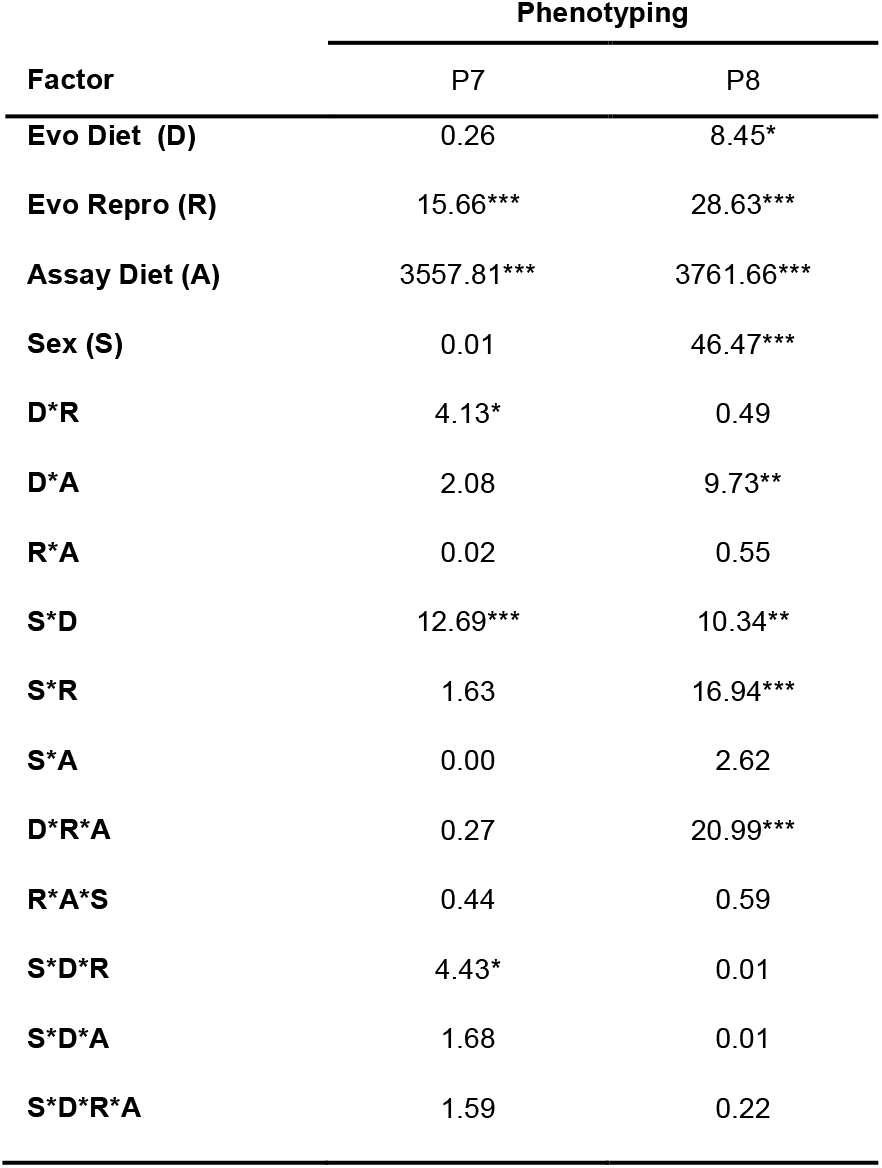
Summary of GLMMs (Chi-square values) for the effect of assay diet (A), evolutionary dietary regime (D) and evolutionary age at reproduction (R) on lifespan across phenotyping sessions. Significance of Chi-square values are indicated by *: * =P<0.05, **=P<0.01, ***=P<0.001.

| Factor | Phenotyping |  |
| --- | --- | --- |
|  | P7 | P8 |
| Evo Diet (D) | 0.26 | 8.45* |
| Evo Repro (R) | 15.66*** | 28.63*** |
| Assay Diet (A) | 3557.81*** | 3761.66*** |
| Sex (S) | 0.01 | 46.47*** |
| D*R | 4.13* | 0.49 |
| D*A | 2.08 | 9.73** |
| R*A | 0.02 | 0.55 |
| S*D | 12.69*** | 10.34** |
| S*R | 1.63 | 16.94*** |
| S*A | 0.00 | 2.62 |
| D*R*A | 0.27 | 20.99*** |
| R*A*S | 0.44 | 0.59 |
| S*D*R | 4.43* | 0.01 |
| S*D*A | 1.68 | 0.01 |
| S*D*R*A | 1.59 | 0.22 |

**Table 4:** Summary of GLMMs (Chi-square values) for the effect of assay diet (A), evolutionary dietary regime (D) and evolutionary age at reproduction (R) on fecundity at early, mid and late ages across phenotyping sessions. Significance of Chi-square values are indicated by *: * =P<0.05, **=P<0.01, ***=P<0.001.

| Phenotyping | Age | Evo Diet (D) | Evo Repro (R) | Assay Diet (A) | D*R | D*A | R*A | D*R*A |
| --- | --- | --- | --- | --- | --- | --- | --- | --- |
| P7 | Early | 1.00 | 1.85 | 176.80*** | 4.45 | 35.26*** | 1.18 | 5.47 |
|  | Mid | 4.69 | 0.49 | 1383.33*** | 8.14* | 8.02* | 20.00*** | 0.72 |
|  | Late | 1.5 | 7.5* | 2154.05*** | 0.7135 | 1.84 | 76.34*** | 7.62* |
| P8 | Early | 0.20 | 1.60 | 892.25*** | 0.77 | 9.05 | 0.49 | 8.05 |
|  | Mid | 0.62 | 0.89 | 364.66*** | 10.20** | 87.46*** | 43.04*** | 132.38*** |
|  | Late | 0.29 | 7.54* | 204.34*** | 0.934 | 84.75*** | 64.578*** | 90.37*** |

However, what hampers firm conclusions about fecundity are the inconsistent phenotypes of the 1.0 line females across the P7 and P8 sessions (Fig. 6). The slightly different assay conditions between the two phenotyping sessions (one male and one female per vial in P7 vs. two females and two males per vial in P8) present one potential cause as females are known to adjust their fecundity based on density (e.g. Barker, 1973; this study compares the difference between vial densities of 5 and 50 females or more). Slight changes in environmental conditions (e.g., note the considerably faster development in P8 relative to P7; Fig. 2) might also have affected overall patterns of fecundity. Clearly, further experiments (e.g., tracking fecundity across the entire reproductive span) are necessary to clarify this issue.

### Does adult body size drive patterns of life history adaptation?

In many studies body size correlates positively with developmental time, lifespan, and fecundity (see above and Robertson, 1957, Hillesheim & Stearns, 1992, Honěk, 1993, Zwaan et al., 1995, Prasad et al., 2001). Our results also showed such correlations; for instance, selection for late life reproduction extended lifespan and increased adult weight for males and females alike. However, these correlations are unlikely to constrain the evolution of the life history adaptations, but will rather modulate them. For instance, while selection for late reproduction consistently increased lifespan for the 0.25 lines, these lines also sped up their development and reduced their weight relative to the 1 and 2.5 lines. Furthermore, the fact that the differences in body weight between early and late life populations were large in early life, but disappeared later in life (at the time of actual selection for the late lines) for 1 and 2.5 but not 0.25 lines (Fig. 7), suggests that body size evolved for a different reason in these lines relative to the 0.25 lines. For instance, increased body size as a response to late life reproduction in the 0.25 lines may serve to increase fecundity in the face of decreased adult weight as an adaptive response to the larval nutritional condition, while in the 1 and 2.5 lines increased body size it may be related to increasing lifespan.

## Conclusions

Our results suggests that adaptation during one life stage may be contingent on the selection pressures experienced in other stages, and that adaptation to two different selection pressures can lead to different life history strategies to those found when adapting to only one selection pressure at a time. In particular, the dependence of lifespan extension on evolutionary developmental diet suggests that developmental acquisition can be an important factor influencing longevity. While there is still relatively little empirical work on adaptation to multiple or opposing selection pressures (but see: Lankau, 2007, Tarwater & Beissinger, 2013), their prevalence in nature means that a better understanding can further our understanding of evolution under natural conditions (reviewed in Schluter et al., 1991). Indeed, the idea that opposing selection pressures constrain trait evolution is one of the hypotheses put forward to explain why, despite strong consistent directional selection on many traits, there is often little change in trait means across generations in natural populations (Merilä et al., 2001, Kingsolver & Diamond, 2011, Siepielski et al., 2011). Given that multiple selection pressures are likely the norm rather than the exception in nature, our findings suggest that trade-offs should be considered not only between traits within an organism, but also between adaptive responses to differing selection pressures.

## Supporting information

## References

Baldal, E. A., Brakefield, P. M. & Zwaan, B. J. 2006. Multitrait evolution in lines of *Drosophila melanogaster* selected for increased starvation resistance: the role of metabolic rate and implications for the evolution of longevity. Evolution 60: 1435–1444.

Barker, J. S. F. 1973. Adult population density, fecundity and productivity in Drosophila melanogaster and Drosophila simulans. Oecologia 11: 83–92.

Bates, D., Maechler, M., Bolker, B. & Walker, S. 2015. Fitting linear mixed-effects models using lme4. Journal of Statistical Software 67: 1–48.

Beldade, P. & Brakefield, P. M. 2002. The genetics and evo-devo of butterfly wing patterns. Nature Reviews Genetics 3: 442–452.

Bochdanovits, Z. & Jong, G. d. 2003. Experimental evolution in *Drosophila melanogaster*: Interaction of temperature and food quality selection regimes. Evolution 57: 1829–1836.

Bubliy, O. A. & Loeschcke, V. 2005. Correlated responses to selection for stress resistance and longevity in a laboratory population of Drosophila melanogaster. Journal of Evolutionary Biology 18: 789–803.

Chippindale, A. K., Chu, T. J. & Rose, M. R. 1996. Complex trade-offs and the evolution of starvation resistance in *Drosophila melanogaster*. Evolution: 753–766.

Clancy, D. J. & Kennington, W. J. 2001. A simple method to achieve consistent larval density in bottle cultures. Drosophila Information Service 84: 168–169.

Davidowitz, G., Roff, D. & Nijhout, H. F. 2016. Synergism and antagonism of proximate mechanisms enable and constrain the response to simultaneous selection on body size and development time: An empirical test using experimental evolution. The American Naturalist 188: 499–520.

de Jong, G. & van Noordwijk, A. J. 1992. Acquisition and allocation of resources: Genetic (co) variances, selection, and life histories. The American Naturalist 139: 749–770.

Economos, A. 1980. Taxonomic differences in the mammalian life span-body weight relationship and the problem of brain weight. Gerontology 26: 90–98.

Fox, J. & Weisberg, S. 2010. An R companion to applied regression. Sage.

Frankino, W. A., Zwaan, B. J., Stern, D. L. & Brakefield, P. M. 2005. Natural selection and developmental constraints in the evolution of allometries. Science 307: 718–720.

Hillesheim, E. & Stearns, S. C. 1992. Correlated responses in life-history traits to artificial selection for body weight in *Drosophila melanogaster*. Evolution 46: 745–752.

Hoffmann, A. A., Hallas, R., Anderson, A. R. & Telonis-Scott, M. 2005. Evidence for a robust sex-specific trade-off between cold resistance and starvation resistance in *Drosophila melanogaster*. Journal of Evolutionary Biology 18: 804–810.

Holm, S. 1979. A simple sequentially rejective multiple test procedure. Scandinavian journal of statistics: 65–70.

Honěk, A. 1993. Intraspecific variation in body size and fecundity in insects: a general relationship. Okios 66.

Kawecki, T. J., Lenski, R. E., Ebert, D., Hollis, B., Olivieri, I. & Whitlock, M. C. 2012. Experimental evolution. Trends in Ecology & Evolution 27: 547–560.

Khazaeli, A. A., Van Voorhies, W. & Curtsinger, J. W. 2005. The relationship between life span and adult body size is highly strain-specific in *Drosophila melanogaster*. Experimental Gerontology 40: 377–385.

Kingsolver, J. G. & Diamond, S. E. 2011. Phenotypic selection in natural populations: what limits directional selection? The American Naturalist 177: 346–357.

Kirkwood, T. B. L. & Holliday, R. 1979. The evolution of ageing and longevity. Proceedings of the Royal Society of London B: Biological Sciences 205: 531–546.

Kirkwood, T. B. L. & Rose, M. R. 1991. Evolution of Senescence: Late Survival Sacrificed for Reproduction. Philosophical Transactions of the Royal Society of London B: Biological Sciences 332: 15–24.

Kolss, M., Vijendravarma, R. K., Schwaller, G. & Kawecki, T. J. 2009. Life-history consequences of adaptation to larval nutritional stress in Drosophila. Evolution 63: 2389–2401.

Kristensen, T. N., Overgaard, J., Loeschcke, V. & Mayntz, D. 2010. Dietary protein content affects evolution for body size, body fat and viability in *Drosophila melanogaster*. Biology Letters.

Lankau, R. A. 2007. Specialist and generalist herbivores exert opposing selection on a chemical defense. New Phytologist 175: 176–184.

Lee, K. P., Simpson, S. J., Clissold, F. J., Brooks, R., Ballard, J. W. O., Taylor, P. W., Soran, N. & Raubenheimer, D. 2008. Lifespan and reproduction in Drosophila: new insights from nutritional geometry. Proceedings of the National Academy of Sciences 105: 2498–2503.

Leftwich, P. T., Nash, W. J., Friend, L. A. & Chapman, T. 2016. Adaptation to divergent larval diets in the medfly, *Ceratitis capitata*. Evolution 71: 289–303.

Leroi, A. M., Chen, W. R. & Rose, M. R. 1994a. Long-term laboratory evolution of a genetic life-history trade-off in *Drosophila melanogaster*. 2. Stability of genetic correlations. Evolution 48: 1258–1268.

Leroi, A. M., Chippindale, A. K. & Rose, M. R. 1994b. Long-term laboratory evolution of a genetic life-history trade-off in *Drosophila melanogaster.* 1. The role of genotype-by-environment interaction. Evolution 48: 1244–1257.

Lints, F. A. 1978. Genetics and ageing. S. Karger.

Luckinbill, L. S., Arking, R., Clare, M. J., Cirocco, W. C. & Buck, S. A. 1984. Selection for delayed senescence in *Drosophila melanogaster*. Evolution 38: 996–1003.

Magwere, T., Chapman, T. & Partridge, L. 2004. Sex Differences in the Effect of Dietary Restriction on Life Span and Mortality Rates in Female and Male Drosophila Melanogaster. The Journals of Gerontology Series A: Biological Sciences and Medical Sciences 59: B3–B9.

May, C. M., Doroszuk, A. & Zwaan, B. J. 2015. The effect of developmental nutrition on life span and fecundity depends on the adult reproductive environment in Drosophila melanogaster. Ecology and Evolution 5: 1156–1168.

Merilä, J., Sheldon, B. & Kruuk, L. 2001. Explaining stasis: microevolutionary studies in natural populations. Genetica 112: 199–222.

Partridge, L. & Fowler, K. 1992. Direct and correlated responses to selection on age at reproduction in *Drosophila melanogaster*. Evolution 46: 76–91.

Prasad, N., Shakara, M., Anitha, D., Rajamani, M. & Joshi, A. 2001. Correlated responses to selection for faster development and early reproduction in Drosophila: the evolution of larval traits. Evolution 55: 1363–1372.

Promislow, D. E. 1993. On size and survival: progress and pitfalls in the allometry of life span. Journal of gerontology 48: B115–B123.

R Development Core Team (2005) A language and environment for statistical computing. pp. R Foundation for Statistical Computing, Vienna, Austria.

Robertson, F. W. 1957. Studies in quantitative inheritance XI. Genetic and environmental correlation between body size and egg production in *Drosophila melanogaster*. Journal of Genetics 55: 428–443.

Roff, D. A. 1992. Evolution of life histories: theory and analysis. Springer Science & Business Media.

Roff, D. A. 2001. Life History Evolution Sinauer Associates, Sunderland.

Rose, M. R. 1984. Laboratory evolution of postponed senescence in *Drosophila melanogaster*. Evolution 38: 1004–1010.

Schluter, D., Price, T. D. & Rowe, L. 1991. Conflicting selection pressures and life history trade-offs. Proceedings of the Royal Society of London. Series B: Biological Sciences 246: 11–17.

Siepielski, A. M., DiBattista, J. D., Evans, J. A. & Carlson, S. M. 2011. Differences in the temporal dynamics of phenotypic selection among fitness components in the wild. Proceedings of the Royal Society of London B: Biological Sciences 278: 1572–1580.

Stearns, S. C. 1992. The Evolution of Life Histories. Oxford University Press, Oxford.

Tarwater, C. E. & Beissinger, S. R. 2013. Opposing selection and environmental variation modify optimal timing of breeding. Proceedings of the National Academy of Sciences 110: 15365–15370.

Therneau, T. M. (2015) coxme: Mixed Effects Cox Models. pp.

Tukey, J. W. 1949. Comparing individual means in the analysis of variance. Biometrics: 99–114.

Van Noordwijk, A. J. & de Jong, G. 1986. Acquisition and allocation of resources: their influence on variation in life history tactics. American Naturalist: 137–142.

Zajitschek, F., Zajitschek, S. R., Canton, C., Georgolopoulos, G., Friberg, U. & Maklakov, A. A. (2016) Evolution under dietary restriction increases male reproductive performance without survival cost. In: Proc. R. Soc. B, Vol. 283. pp. 20152726.

The Royal Society. Zwaan, B., Bijlsma, R. & Hoekstra, R. 1995. Artificial selection for developmental time in *Drosophila melanogaster* in relation to the evolution of aging: direct and correlated responses. Evolution: 635–648.

Zwaan, B. J. 1999. The evolutionary genetics of ageing and longevity. Heredity 82: 589–597.

Zwaan, B. J., Bijlsma, R. & Hoekstra, R. F. 1991. On the developmental theory of ageing. I. Starvation resistance and longevity in *Drosophila melanogaster* in relation to pre-adult breeding conditions. Heredity 66: 29–39.

