## Supplementary material for "Adaptation to developmental diet influences the response to selection on age at reproduction in the fruit fly"

Supplementary Table 1: Diet composition per liter of water

| Diet composition | 0.25 | 1.0 | 2.5 |
|---|---|---|---|
| **Yeast*** | 17.5g | 70g | 175g |
| **Sugar †** | 25g | 100g | 250g |
| **Agar** | 20g | 20g | 20g |
| **Nipagin solution** | 15mL | 15mL | 15mL |
| **Propionic acid** | 3mL | 3mL | 3mL |

*Fermipan Red Label instant yeast
†Suiker Unie Granulated Sugar Extra Fine

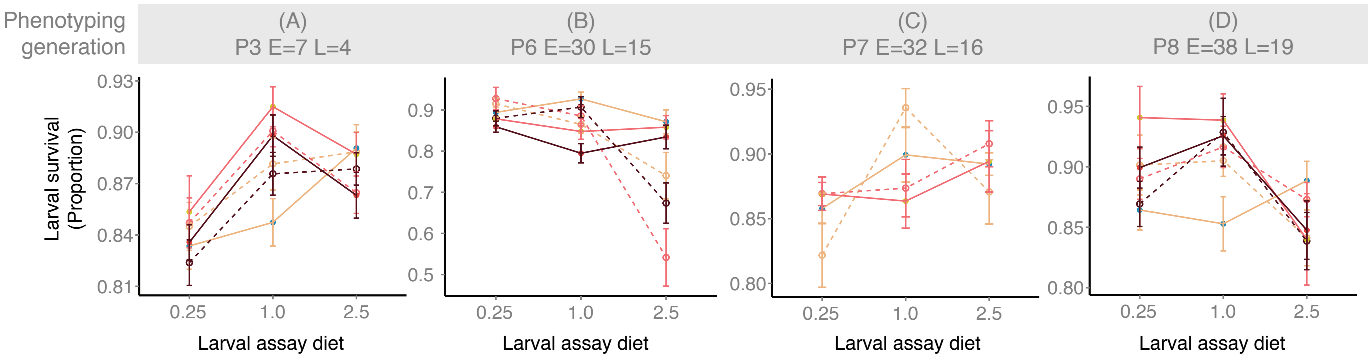


**Supplementary Figure 1: Larval survival from egg to adult (y-axis) across assay diets (x-axis) and phenotyping sessions for each combination of evolutionary dietary regime (colour) and age at reproduction (line type).** Beige, pink, and purple represent the 0.25, 1.0 and 2.5 EE larval diets respectively. Solid and dashed lines represent early (E) and late (L) reproducing lines respectively. All error bars are standard errors of the mean across replicate lines. For simplicity, only phenotyping sessions carried out using all three assay diets are included: P3 (a), P6 (b), P7 (c) and P8 (d). Overall, larval survival was very high across assay diets, ranging from 80 to 95% survival in all but one phenotyping session (P6, b.). Furthermore, the plastic effect of assay diet on CE lines was insignificant in all but one (P5) out of five phenotyping sessions (Table 2). Given the lack of an effect of larval assay diet on survival in the control lines (CE), it is perhaps not surprising that there was also no consistent evolved response. Across all eight phenotyping sessions we detected significant effects of evolutionary regime on larval survival in only three of them (P5, P7, and P8) however, the factors playing a significant role did not overlap (Table 1). For example in P6, L populations displayed a clear reduction in survival relative to E populations under 2.5 assay conditions (Reproductive regime x Assay interaction: $\chi 2=39.86$, $p=<0.0001$; Supplementary Figure 1b), however, this response disappeared in both subsequent phenotyping sessions (Table 1, Supplementary Figure 1c, d). The same pattern is true of all other sessions – while significant effects arise, they are not consistent across generations (Table 1). Thus it appears that while larval survival does appear to be sensitive to transient effects, there is no consistent response of larval survival to the two selection regimes or their interaction. Previous experiments subjecting flies to evolution on very poor larval diets did find increased larval survival in the course of evolution (Kolss, Vijendravarma et al. 2009). However, we deliberately chose our larval diets in such a way that they did not affect viability. In contrast, the diet imposed by Kolss et al., (2009) contained considerably less nutrients than our 0.25 diet, and decreased larval survival by 20% in control lines. Since we observed little difference in larval survival of 1-E lines across assay diets (Supplementary Fig. 1) it is likely that there was very little direct selection for increased larval survival in our experiment. While it is still possible that changes in larval survival could have evolved as a correlated response to other aspects of selection (e.g. in a trade-off with faster larval development or longer lifespan) our results do not support this hypothesis.
